# New Evidence of Altered Gut Microbiota in Autism Spectrum Disorders through Metagenomic Analysis

**DOI:** 10.1101/2025.09.11.675541

**Authors:** Peiqi Xing, Zhaohua Li, Zhuang Xiong, Yingke Ma, Menglin Sun, Yiming Bao

**Affiliations:** China National Center for Bioinformation, Beijing, China, Beijing 100101, China; Beijing Institute of Genomics, Chinese Academy of Sciences, Beijing 100101, China; National Genomics Data Center, China National Center for Bioinformation, Beijing 100101, China; CAS Key Laboratory of Genomic and Precision Medicine, Beijing Institute of Genomics, Chinese Academy of Sciences and China National Center for Bioinformation, Beijing 100101, China; University of Chinese Academy of Sciences, Beijing 100049, China; Wucailu Center for Children with Autism, Wucailu Research Institute, Beijing 100029, China

**Keywords:** Autism Spectrum Disorders, Gut Microbiota, Shotgun Metagenome, Brain-gut Axis, Longitudinal Study

## Abstract

Cumulative evidence suggests alterations of gut microbiota in children with Autism Spectrum Disorder (ASD). However, due to highly diverse research approaches, the distinctive microbial patterns and underlying mechanisms driving ASD etiology remain elusive. Here, we constructed the metagenomic dataset of gut microbiota derived from Chinese children with ASD, incorporating a longitudinal study to explore the pivotal role of gut microbiota in the pathogenesis of ASD. At the bacterial species level, we identified a notable increase in the relative abundance of *Clostridium scindens*, which exhibited a positive correlation with the cholate degradation pathway and showing a decline concurrent with improved behavior in ASD children. Within the fungal community, we observed significant dysbiosis in twenty-six species, underlining their robust discriminatory potential between ASD and neurotypical subjects. Metabolic pathways related to mitochondrial dysfunction were enriched in ASD, alongside peptidoglycan, glutamate, and nucleotide pathways, suggesting potential interactions between the gut and brain through the innate immune system and vagus nerve in ASD. Our findings underscore the significance of the fungal community and mitochondrial dysfunction in ASD etiology, while also shedding light on how perturbed metabolic functions of gut microbiota collectively contribute to the pathogenesis of ASD.

## Introduction

Autism spectrum disorders (ASD) refers to a range of neurodevelopmental disorders, primarily characterized by deficiency in social interactions, communication, and stereotypic or repetitive behaviors. The prevalence of ASD in children and adolescents is 1 in 54 in the United States [1]. It is widely recognized that both genetic and environmental factors are related to the etiology of ASD [2]. Besides the core symptoms, evidence shows that gastrointestinal (GI) disorders and associated symptoms, including constipation, abdominal pain, diarrhea, gaseousness and vomiting, are also reported in individuals with ASD [3]. The GI microbiota influences brain development and behavior through neuroendocrine, neuroimmune and autonomic nervous systems [4]. Findings in this area formed the brain-gut axis concept, a bidirectional communication network between the brain and gut in both the enteric nervous system (ENS) and the central nervous system (CNS).

Based on the concept of the brain-gut axis, the gut microbiota has been suggested to play a role in ASD. Studies conducted on animal models of ASD highlight gut-brain connections that contribute to the pathophysiology of ASD [5–8]. Maternal immune activation (MIA) mouse model that with GI barrier defects and microbiota alterations exhibit features of ASD. Oral treatment of MIA offspring with the human commensal *Bacteroides fragilis* corrects gut permeability, alters the microbial composition, and ameliorates defects in communicative, stereotypic, anxiety-like and sensorimotor behaviors [5]. The gut microbiota transplanted from human donors with ASD regulates behaviors in mice via the production of neuroactive metabolites [6]. However, the results of human studies seem to be unclear. Most of human studies have demonstrated study-specific aberrations in both gut microbiota and their metabolites in ASD (as summarized in Table1 and 2). Species at different taxonomy levels are involved in microbial dysbiosis [9–15] or no significant difference [16, 17] between ASD and TD individuals are observed. Although the role of intestinal flora in the pathogenesis of autism is evident, the consensus across studies based on stool samples from individuals with ASD has not yet been established. This is due to small sample size that is prone to bias (e.g., n<50) [18, 19], inconsistent statistical analysis [9, 18, 20] and different races or geographic environments that dominates in shaping human gut microbiota [21] (Table 1). Furthermore, there are still technical limitations: most studies have used 16S rRNA sequencing (with the exception of a few [22, 23]), which provides limited taxonomic resolution and functional information about the microbiome.

**Table 1.** Summary of relevant literature in the past ten years - research strategy. **Notes:** bTEFAP, bacterial tag-encoded FLX amplicon pyrosequencing; qPCR, quantitative PCR.

| Research | country | case : sibling : control | Method |
| --- | --- | --- | --- |
| Finegold et al., 2010[9] | USA | 33:07:08 | bTEFAP |
| Wang et al., 2011[83] | Australia | 23:22:09 | qPCR |
| Gondalia et al., 2012[84] | Australia | 51:53:00 | bTEFAP |
| De Angelis et al., 2013[18] | Italy | 10:00:10 | bTEFAP & 16S rRNA (active bacteria) & 16S rDNA (total bacteria) |
| Wang et al., 2013[33] | Australia | 23:22:09 | qPCR |
| Kang et al., 2013[10] | USA | 20:00:20 | bTEFAP & 16S rDNA (V2-V3) |
| Tomova et al., 2015[16] | Slovakia | 10:09:10 | qPCR |
| Son et al., 2015[27] | USA | 59:44:00 | qPCR & 16S rRNA(V1-V2&V2-V3) |
| Strati et al., 2017[11] | Italy | 40:00:40 | 16S rRNA (V3-V5) & fungal ITS1 region |
| Berding et al., 2018[85] | USA | 26:00:32 | qPCR & 16S rRNA (V3-V4) |
| Coretti et al., 2018[86] | Italy | 11:00:14 | 16S rRNA (V3-V4) |
| Kang et al., 2018[19] | USA | 21:00:23 | 454 16S rRNA |
| Pulikkan et al., 2018[36] | India | 30:00:24 | 16s rRNA (V3) |
| Rose et al., 2018[87] | USA | 50:00:41 | 16S rRNA (V3-V4) |
| Zhang et al., 2018[14] | China | 35:00:06 | 16S rRNA (V3-V4) |
| Li et al., 2019[88] | China | 59:00:30<br>(mother-child<br>pairs) | 16s rRNA (V1-V2) |
| Liu et al., 2019[13] | China | 30:00:20 | 16s rRNA (V3-V4) +fecal<br>SCFAs |
| Ma et al., 2019[12] | China | 45:00:45 | 16s rRNA (V3-V4) |
| Plaza-Diaz et al., 2019[28] | Spain | 57:00:57 | 16S rRNA (V3-V4) |
| Sun et al., 2019[89] | China | 9:00:06 | 16s rRNA |
| Wang et al., 2019[23] | China | 92:00:42 | Shotgun metagenome |
| Zhai et al., 2019[90] | China | 78:00:58 | 16S rRNA (V3-V4) |
| Ding et al., 2020[34] | China | 77:00:50 | 16s rRNA |
| Dan et al., 2020[22] | China | 143:00:143 | 16S rRNA (V3-V4) + Shotgun<br>metagenome (30 vs 30) |
| Zhang et al., 2020[29] | China | 39:00:40 | Shotgun metagenome |
| Mingxing et al., 2021[91], | China | 773:00:429 | 16s rRNA |

In this study, we employed a shotgun metagenomics sequencing strategy on the largest combined Chinese cohort to confirm the dysbiosis of gut microbiota between children with ASD and typically developing children, and to investigate the microbiota’s functional contributions to the etiology of ASD. Furthermore, we selected five ASD children whose behavior showed improvement following intervention for secondary stool collection and subsequent metagenome sequencing. Additionally, we investigated the potential implications of the identified results on nucleoside, short fatty acid, amino acid, and neurotransmitter metabolism, and elucidated their relevance to ASD etiology.

## Results

### Subject characteristics and study design

In the present study, we enrolled 50 children diagnosed with ASD from the Wucailu Child Behavioral Intervention Center and 14 TD children from nursery and primary schools in Beijing, China. The diagnosis of ASD was based on Autism Diagnostic Observation Schedule (ADOS) evaluation, which was a current ‘gold standard’ diagnostic tool. Considering the limitations of the sample size of our enrolled cohort, we combined age and ethnic matched public dataset that consisted of 43 ASD children and 31 TD children [23]. Table 1 shows the general characteristics of participants. Overall, there is a total of 93 children with ASD and 42 TD children in our combined cohort.

Fifty children with ASD and 14 TD children in our cohort underwent stool sample collection as the first batch. Subsequently, we conducted a year-and-a-half-long follow-up specifically focused on the recruited ASD children. Five children whose behavior improved after the Applied Behavior Analysis (ABA) intervention underwent secondary stool collection (see Method). All stool samples were subjected to shotgun metagenome sequencing. These data, along with a public metagenome dataset (PRJEB23052), were analyzed using the same analytical pipeline to discover the difference between ASD and TD.

The final raw read data were processed at the taxonomic level using MetaPhlAn2 [24] and Kraken2 [25], at functional level using HUMAnN2 [26] (Fig. 1A-D).

**Figure 1.**
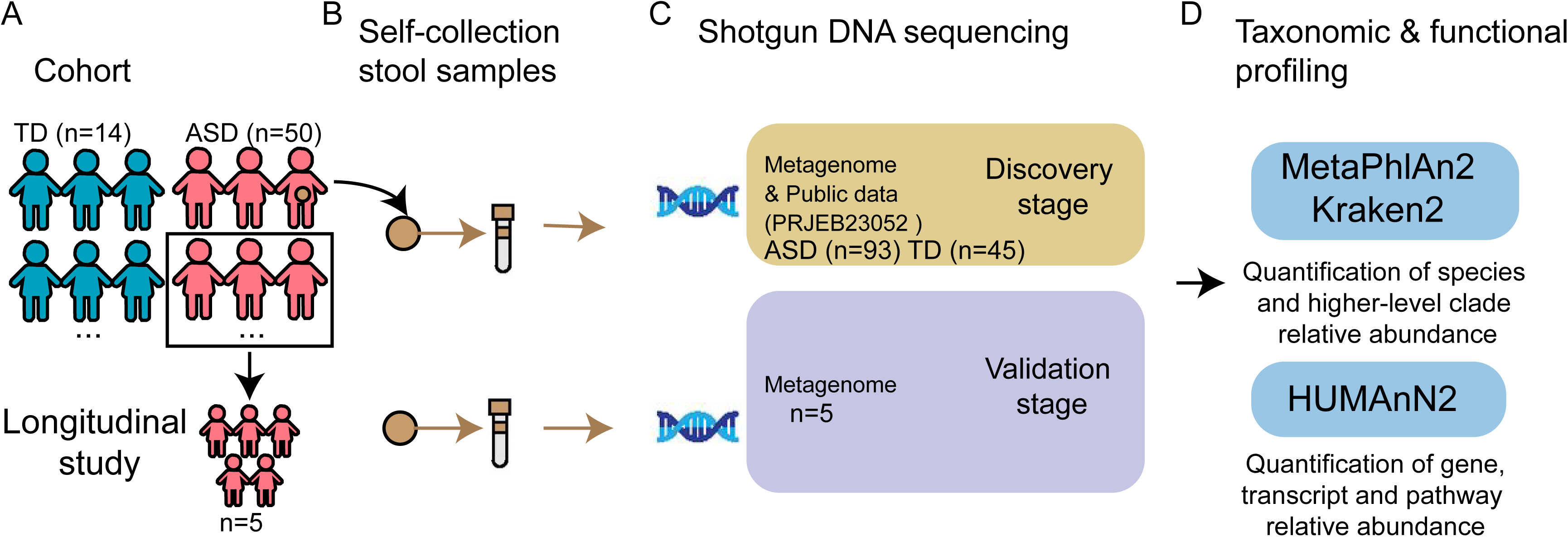
Study design. (A) 50 children with ASD and 14 TD children were enrolled in this study and their stool samples were collected. As a follow-up, we chose 5 children whose behavior improved after intervention to perform secondary stool collection (B) Subjects self-collected samples of stool, which were returned to the laboratory and frozen at −80 °C until DNA extraction. (C) In the discovery stage, the metagenomes of the first batch of samples from all enrolled children were sequenced. These data, combined with a public dataset, were analyzed to discover the difference between ASD and TD. In the validation stage, the metagenomes of 5 follow-up samples were sequenced. (D) Taxonomic relative abundance was evaluated using MetaPhlAn2 and Kraken2, while functional quantification was assessed using HUMAnN2.

### Autistic subjects harbored an altered bacterial gut microbiome

We investigated the gut microbiota of our study cohort through shotgun metagenomics sequencing. Following quality control, an average of 25,712,760 (range 17,265,809– 40,069,242) reads per sample (Table S1) were obtained. The non-metric multidimensional scaling (NMDS) analysis showed the minimal differences between ASD and TD (Fig. 2A, Adonis test: R² = 0.021, *p* < 0.05). We did not observe significant differences in Shannon diversity consistent with previous studies [27–29] (Fig. S1A). The assessment of microbial composition of the ASD and control groups at the phylum level (Fig. 2B) showed top 5 phyla, including *Firmicutes, Bacteroidetes, Actinobacteria, Proteobacteria* and *Verrucomicrobia* constituted the majority of the gut microbiota. The ratio of *Firmicutes/Bacteroidetes* was also not significantly different between the ASD and TD children (Fig. S1B), which is consistent with previous Chinese cohort studies [12, 15]. Genus-level bacteria composition showed great metabolic diversity among samples (Fig. 2C). As enterotype is used to describe the gut microbial community landscape and relevant in clinical practice [30], we conducted enterotype analysis based on the genus level bacteria to figure out if ASD influences enterotype in children. Although there is no significant difference in the percentage of three distinct enterotypes (P/F/B) between ASD and TD, we found more proportion of B enterotype in TD than ASD and the P enterotype was the minimal proportion in the two groups (Fig. S1C). To further identify specific bacteria that are associated with ASD, after linear discriminant effect size (LEfSe) [31] (the linear discriminant analysis (LDA) score >2) and Wilcoxon rank-sum test (adjusted *p*-value < 0.05), we found significant different bacteria taxon between ASD and TD (Fig. 2D-E, Table S2). At the genus level, 8 genera showed significant differences, with *Eggerthella* and *Lachnoclostridium* enriched in ASD and *Sutterella* enriched in TD. These findings are consistent with previous studies [23, 32–34]. At the species level, we identified 14 species which were mostly study-specific in other related studies.

**Figure 2.**
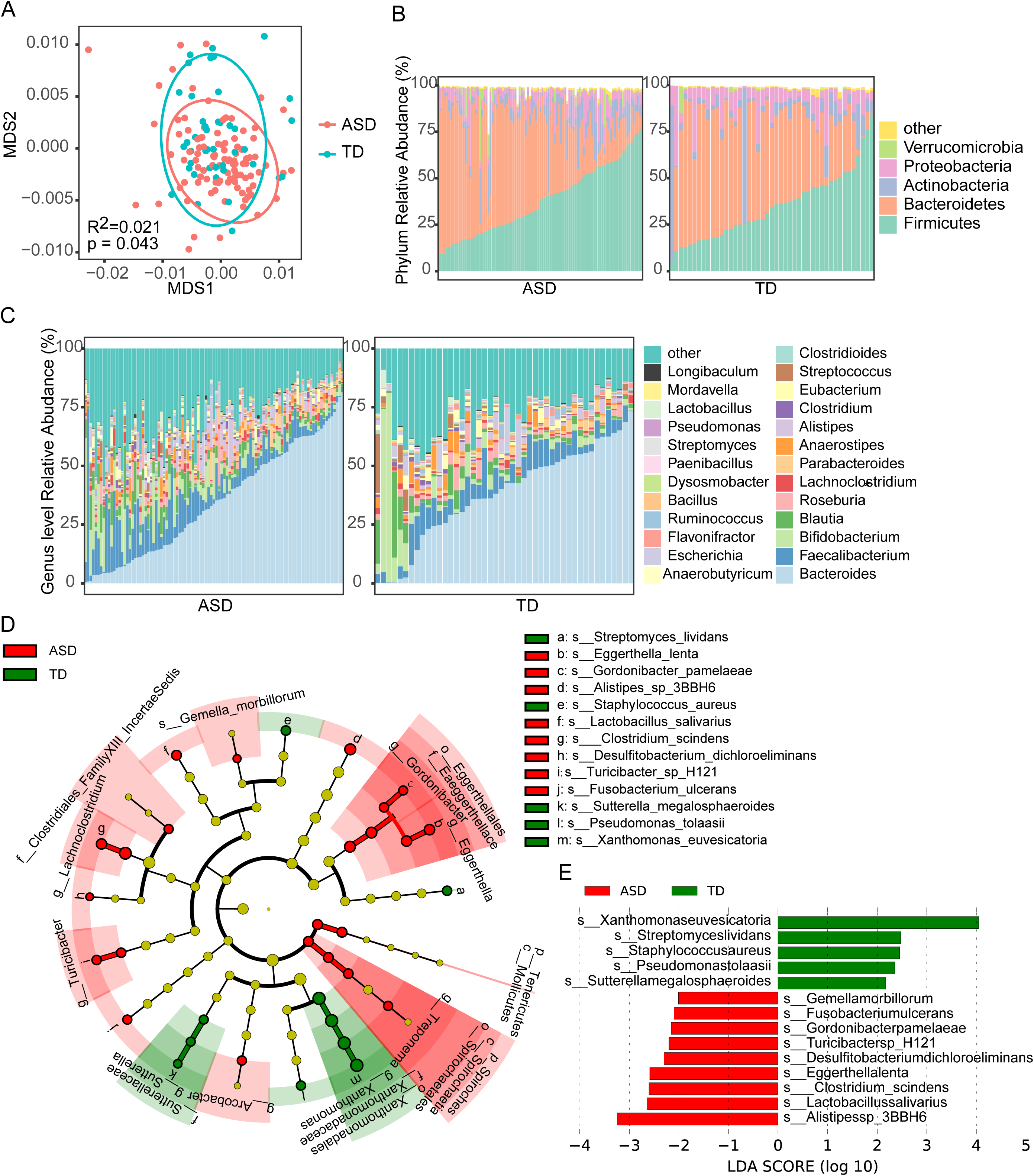
Autistic subjects harbored an altered bacterial gut microbiome. (A) Non-metric Multidimensional Scaling (NMDS) of the Bray–Curtis distance for the species level gut bacterial communities. Ellipses represent 90% CI. R^2^ represents the proportion of variation explained by the NMDS analysis, and *p*-values were calculated by Adonis. (B) Relative abundance of bacterial phyla in the intestinal microbiota. (C) Relative abundance of bacterial genera in the intestinal microbiota. (D) Cladograms generated by Linear discriminant analysis Effect Size (LEfSe) indicating differences in the bacterial taxa (from phylum to species) between TD and ASD. Red nodes indicate taxa that were enriched in ASD, green nodes indicate taxa that were enriched in TD. (E) LDA scores for the species differentially abundant between TD and ASD (LDA > 2). Red bars indicate taxa that were enriched in ASD, and green bars indicate taxa that were enriched in TD. Values that are significantly different by the Wilcoxon rank sum test are indicated by asterisks as follows: *, adj.*p* < 0.05; **, adj.*p* < 0.01; ***, adj.*p* < 0.001.

Pieces of evidence link the colonization by the bacteria genus of *Clostridium* with neurological symptoms and/or etiology of ASD in human individuals [35]. Although our results did not show a significant difference in *Clostridium* abundance, we identified 5 species from *Clostridium*, namely *C. botulinum, C. cellulosi, C. sphenoides, C. hiranonis and C. scindens,* which were significantly elevated in ASD (Wilcoxon rank-sum test, p < 0.01) (Fig. 3A-E). And notably, the relative abundance of *C. scindens* showed a consistent decrease as behavior improved in our follow-up samples (Fig. 3F). Combining comparison results from MetaPhlAn2 (Table S3), we also identified 4 bacterial strains, *C. scindens* GCF000154505*, C. bartlettii GCF000154445, Eggerthella lenta GCF000024265* and *Lachnospiraceae bacterium-5-1-57FAA GCF000218425,* that were significantly increased in the ASD group (Fig. S2A-D) and decreased in the follow-up samples as behavior improved (Fig. S2E-H).

**Figure 3.**
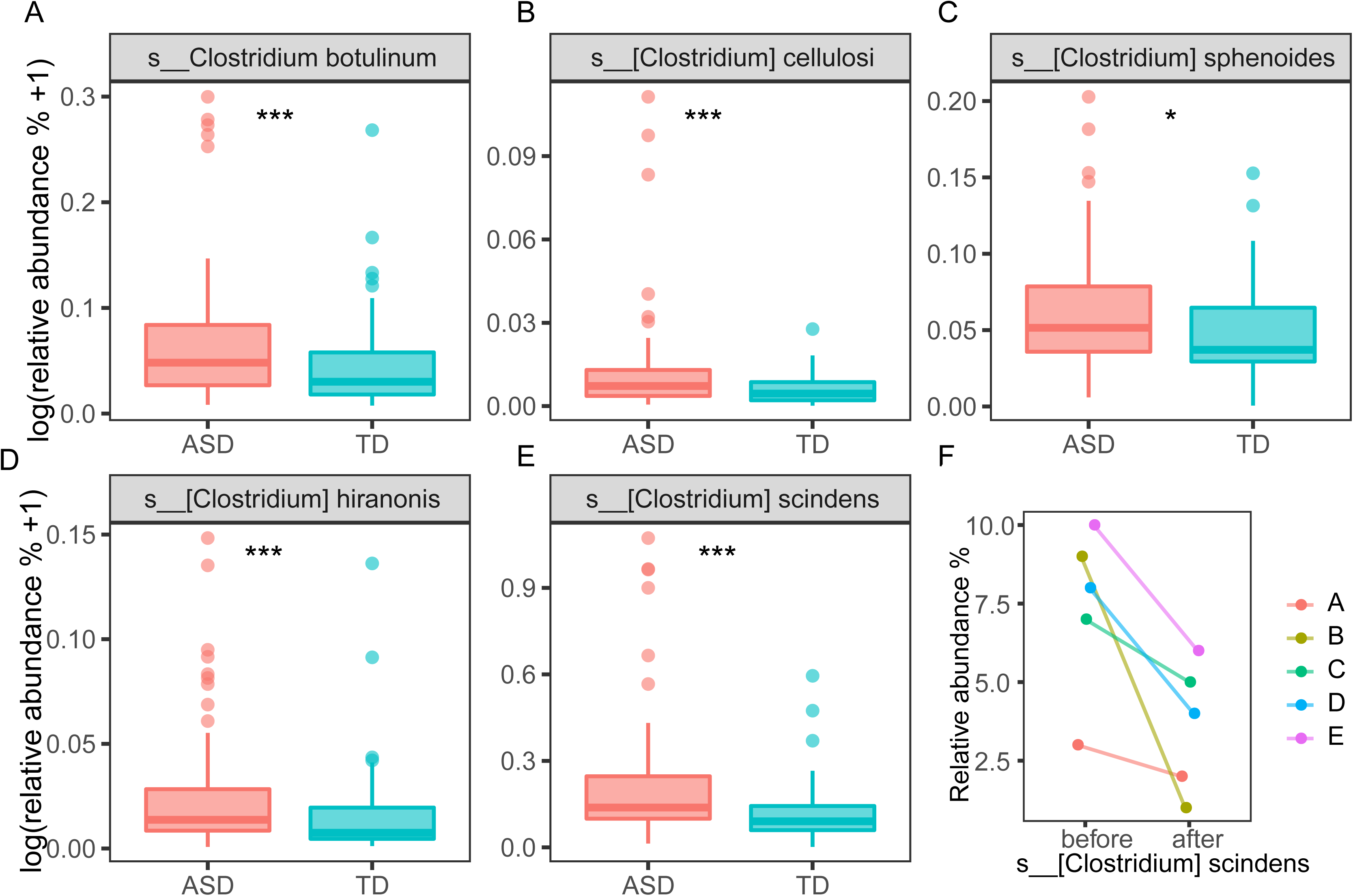
Comparison of the relative abundance of clostridium species between ASD and TD. (A-E) Relative abundance of *Clostridium botulinum*, *Clostridium cellulosi*, *Clostridium sphenoides*, *Clostridium hiranonis* and *Clostridium scindens.* (F) Variation of relative abundance of *Clostridium scindens* in 5 follow-up ASD children (A/B/C/D/E). Values that are significantly different by the Wilcoxon rank sum test are indicated by asterisks as follows: *, *P* < 0.05; **, *P* < 0.01; ***, *P* < 0.001.

### Distinct dysbiosis in fungi community found in autistic subjects

Although the relative abundance of eukaryotic fungi did not show significant differences between the ASD and TD groups (Fig. 4A), we observed a clear separation based on NMDS (Fig. 4B) and significant differences in both Shannon and Simpson diversity indices between the two groups (Fig. 4C). The top 15 genera of fungi composition showed great microbiome diversity among samples (Fig. 4D). We also identified 18 genera that showed a remarkable difference, among which only one genus, *Fusarium*, showed a decreased in ASD (Table S4). Twenty-six species in the fungi community that with significant dysbiosis showed obvious separation between groups (Table S4, Fig. 4E). Three species, *C. albicans, C. dubliniensis* and *C. glabrata*, from *Candida* consistently increased in ASD which not only further confirmed the potential role of *Candida* dysbiosis in ASD but also pinpointed to specific species (Fig. 4F). These results suggested that the fungi community might have a stronger degree of discrimination for ASD.

**Figure 4.**
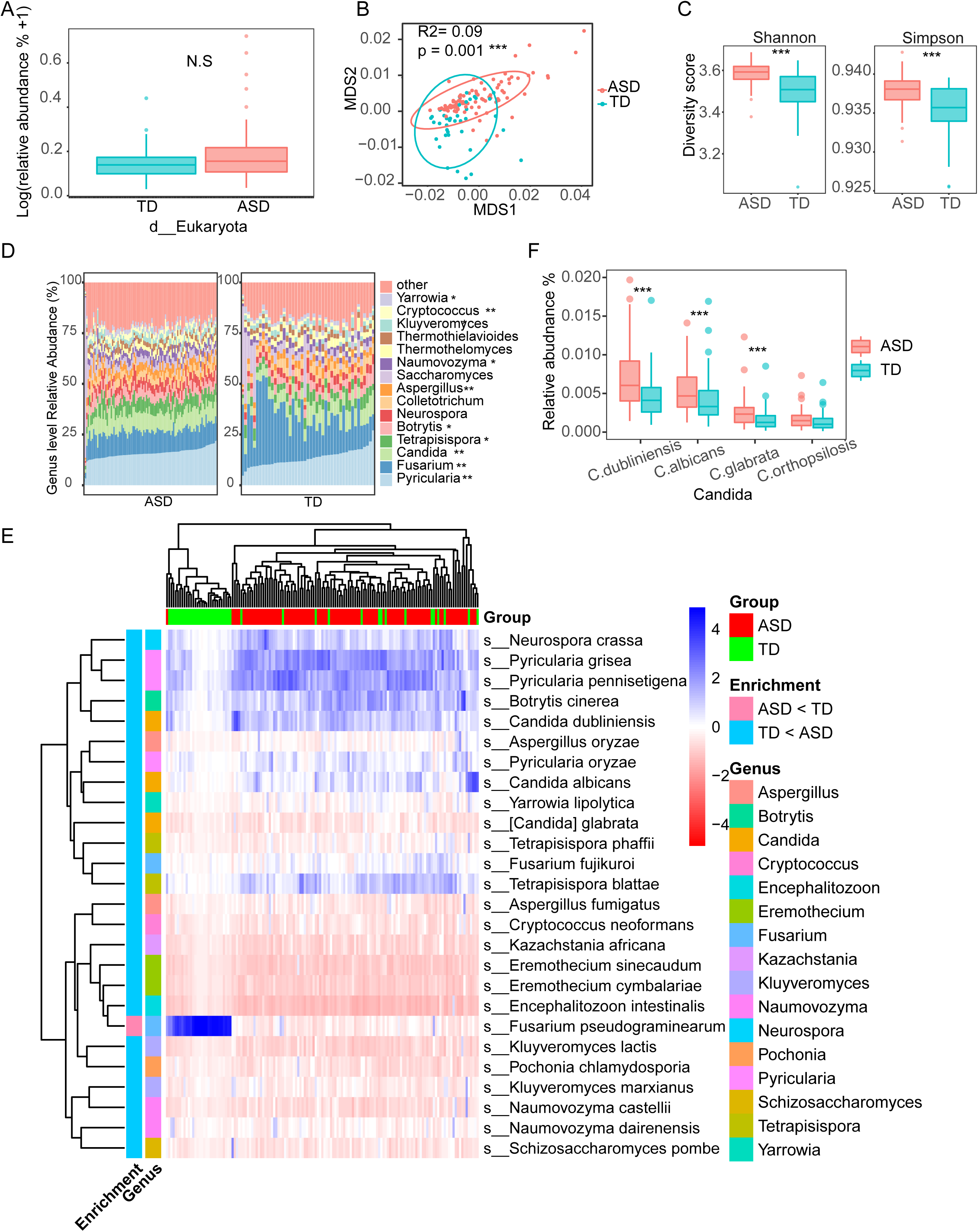
Distinct dysbiosis in fungi community between ASD and TD. (A) Relative abundance of gut eukaryotes between the ASD and TD groups. (Wilcoxon rank sum test, *P* > 0.05) (B) Non-metric Multidimensional Scaling (NMDS) of the Bray–Curtis distance for the species level of gut fungi communities. Ellipses represent 90% CI. R^2^ and *p* value were calculated by Adonis. (C) Shannon and Simpson diversities of fungi community were independently calculated for the ASD and TD groups. (Wilcoxon rank sum test, *P* < 0.01) (D) Relative abundance of fungi genera in the intestinal microbiota. (E) Heatmap of relative abundance of significantly different fungi species. (F) Comparison of relative abundance of three species from *Candida*.

### Diagnostic model based on species level of gut microbiota

In order to corroborate the predictive role of gut microbiota in ASD, we constructed a random forest classifier. Based on the relative abundance of 318 species of gut microbiota, ROC evaluation with 1000 bootstrap showed that the model accurately described the deviations between ASD and control subjects and achieved a diagnostic power as high as 92.59% AUC (Fig. 5A, AUC1). The diagnostic model also provides the contribution of each species to the model. Cross-validation of the contributions of all species showed that selecting the top 43 species yielded the lowest error rate (Fig. 5B). Based on these 43 species we also get a higher AUC (Fig. 5A, AUC2). The 30 species with the highest contribution to the model were shown in Fig. S3. From this list of top species, we found not only *Candida dubliniensis* and *Pyricularia grisea*, which were significantly dysregulated in fungal populations, but also multiple species from *Bifidobacterium* and *Lactobacillus*, such as *B. pseudolongum*, *B. longum*, *B. animalis*, *B. adolescentis*, *L. fermentum*, *L. mucosae*, *L. plantarum* and *L. salivarius* (Fig. S3). Among them, *Bifidobacterium* has been repeatedly reported to be reduced in the gut of ASD patients [9, 18, 29, 36], while *Lactobacillus* spp has been repeatedly observed to be elevated in the gut of ASD patients [11, 16, 29, 36] (as summarized in Table 2). It is also worth noting that multiple species of these two genera are used as probiotics to intervene ASD children and have certain therapeutic effects [37–39], including *B. longum, L. fermentum, L. plantarum, L. acidophilus,* and *L. rhamnosus*. Therefore, we also paid attention to the distribution of these two genera in the cohort. As shown in Fig. 5C, several *Lactobacillus* species showed extremely high individual differences in the ASD group, and the content was significantly increased in specific individuals. The other four *Bifidobacterium* species showed similar trends to *Lactobacillus* species except for *B. longum* (Fig. 5D), which as a probiotic for ASD intervention. The strong individual heterogeneity of this intestinal microbiota in children with ASD not only explain the inconsistent findings of gut microbiota dysbiosis identified in different studies, but also further underscores the precision probiotic intervention and the identification of differential strains in the gut microbiota of ASD.

**Figure 5.**
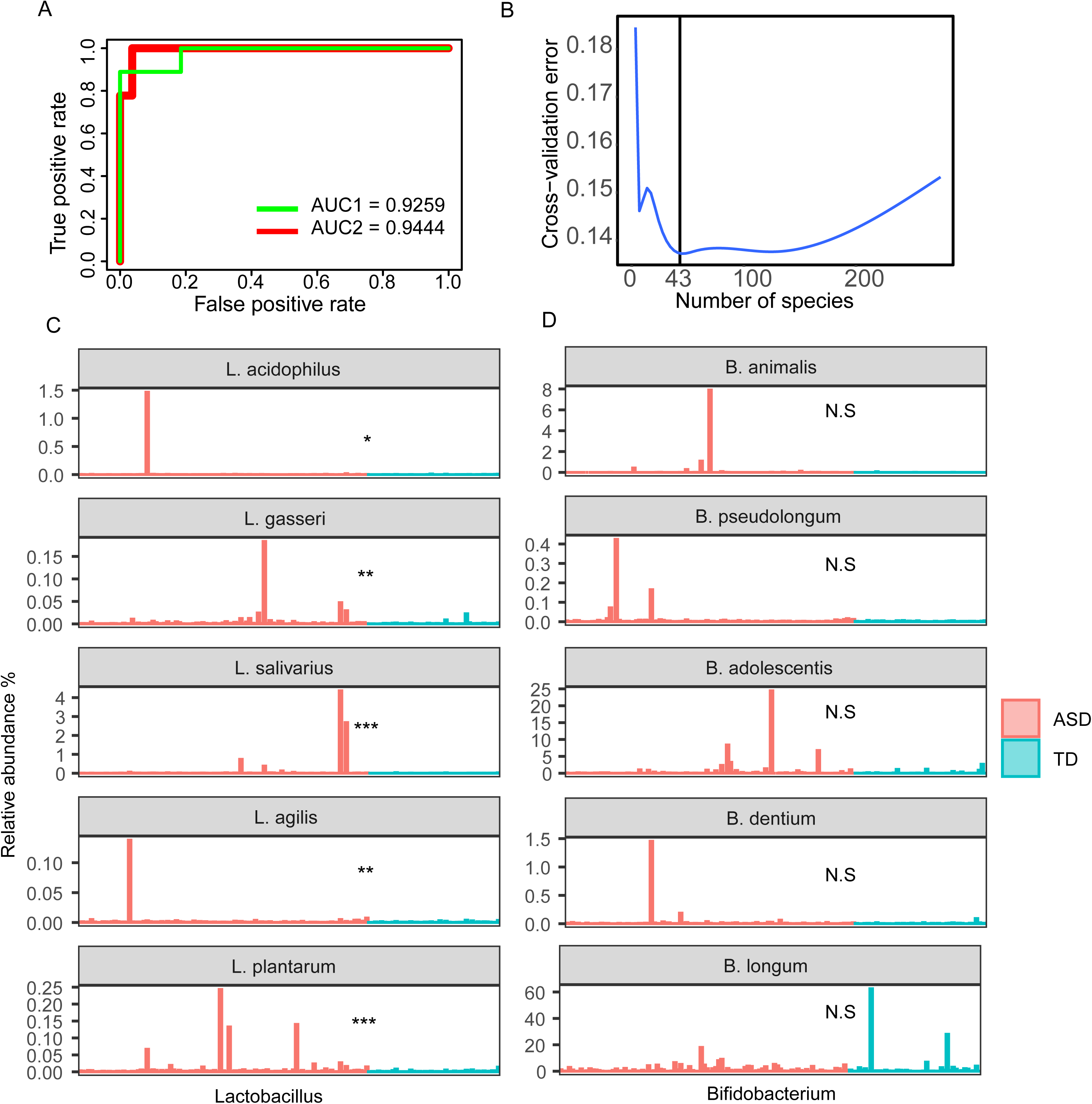
The performance of random forest model based on gut microbiota species between ASD and TD. (A) ROC analysis of the performance of the diagnostic model. Green line indicate model based all species, red line indicate model based top 43 contributed species. Values that are significantly different by the Wilcoxon rank sum test are indicated by asterisks as follows: *, adj.*p* < 0.05; **, adj.*p* < 0.01; ***, adj.*p* < 0.001. (B) Error rate distributions species selection through cross validation. (C-D) The longitudinal directions of figures C and D represent the distribution of 5 species belonging to the genus of *Lactobacillus* and *Bifidobacterium* respectively between children with or without ASD. Values that are significantly different by the Wilcoxon rank sum test are indicated by asterisks as follows: *, *p* < 0.05; **, *p* < 0.01; ***, *p* < 0.001.

**Table 2.**
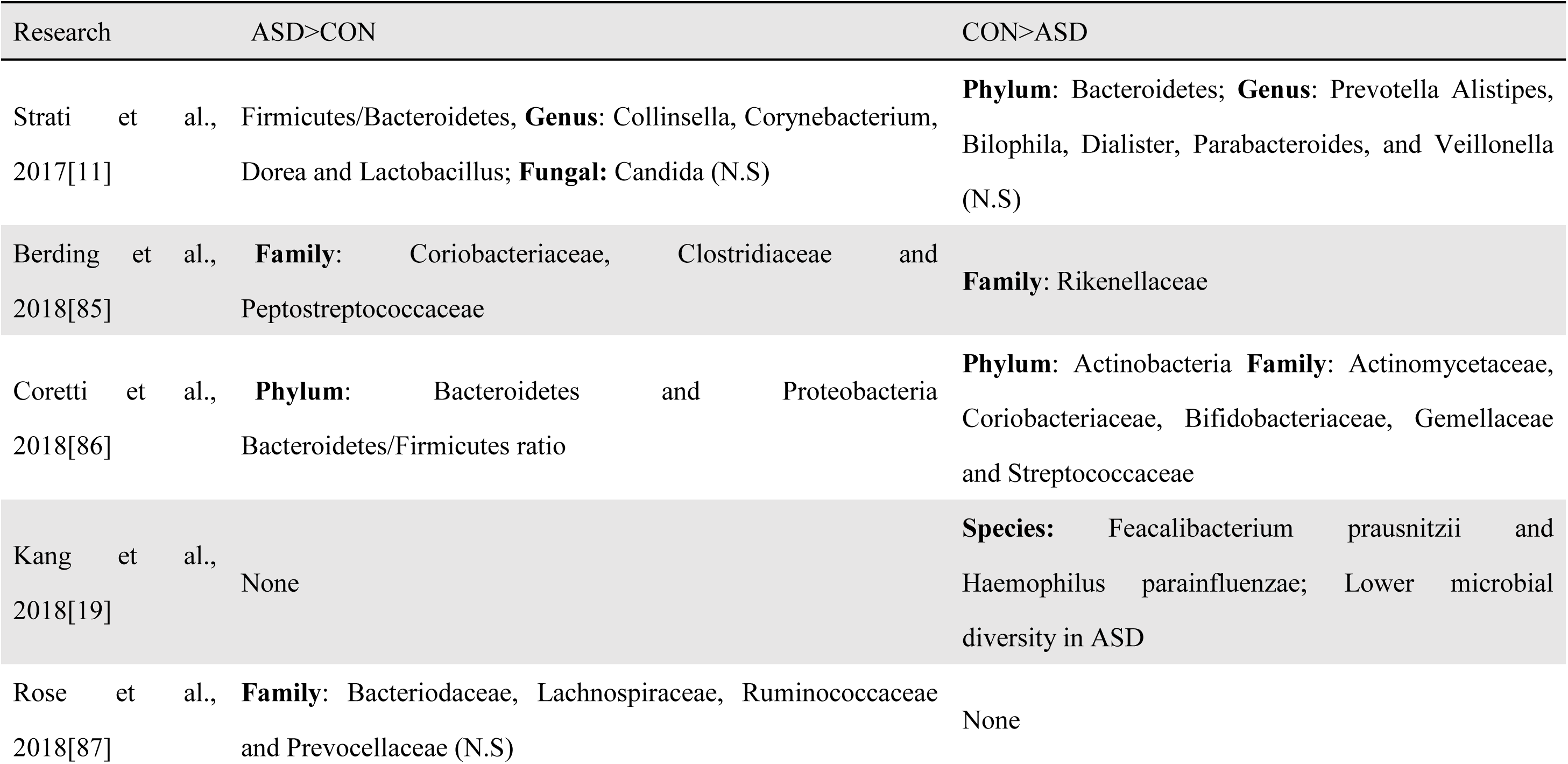
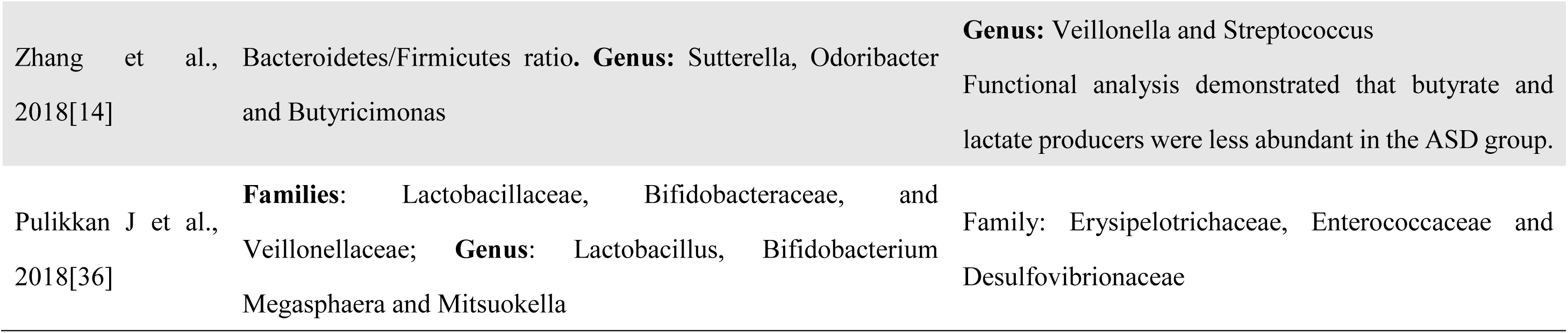
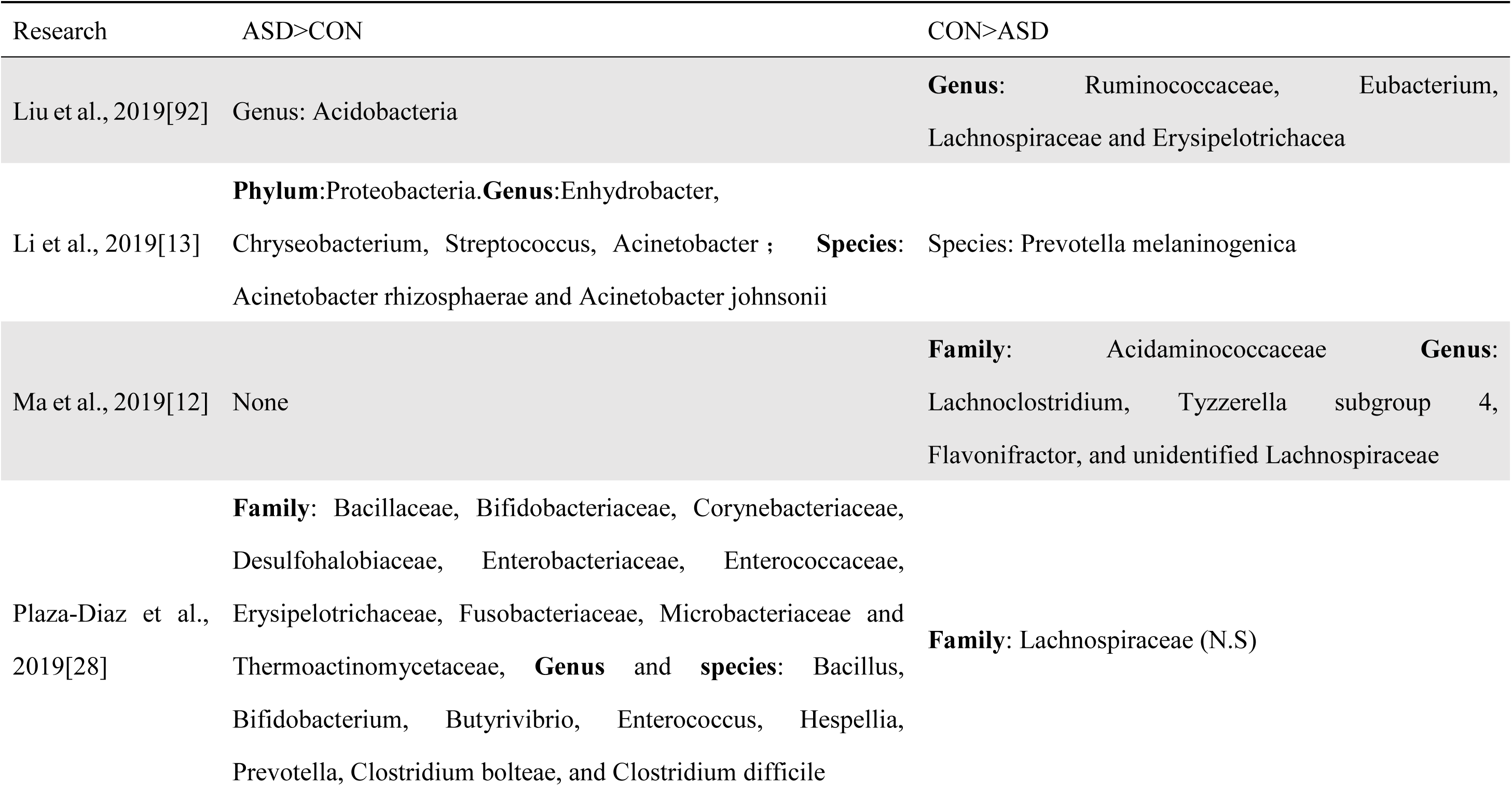
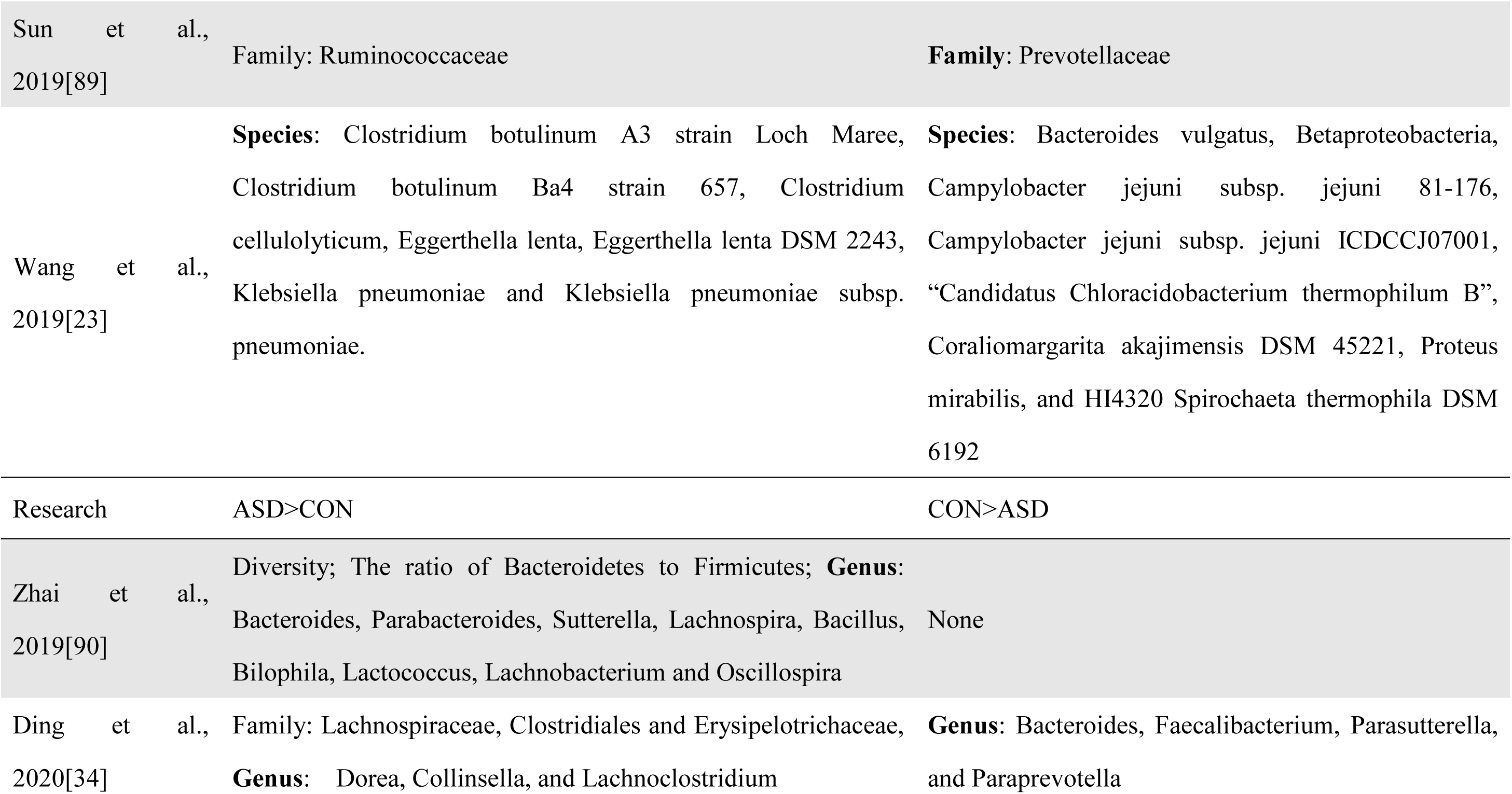
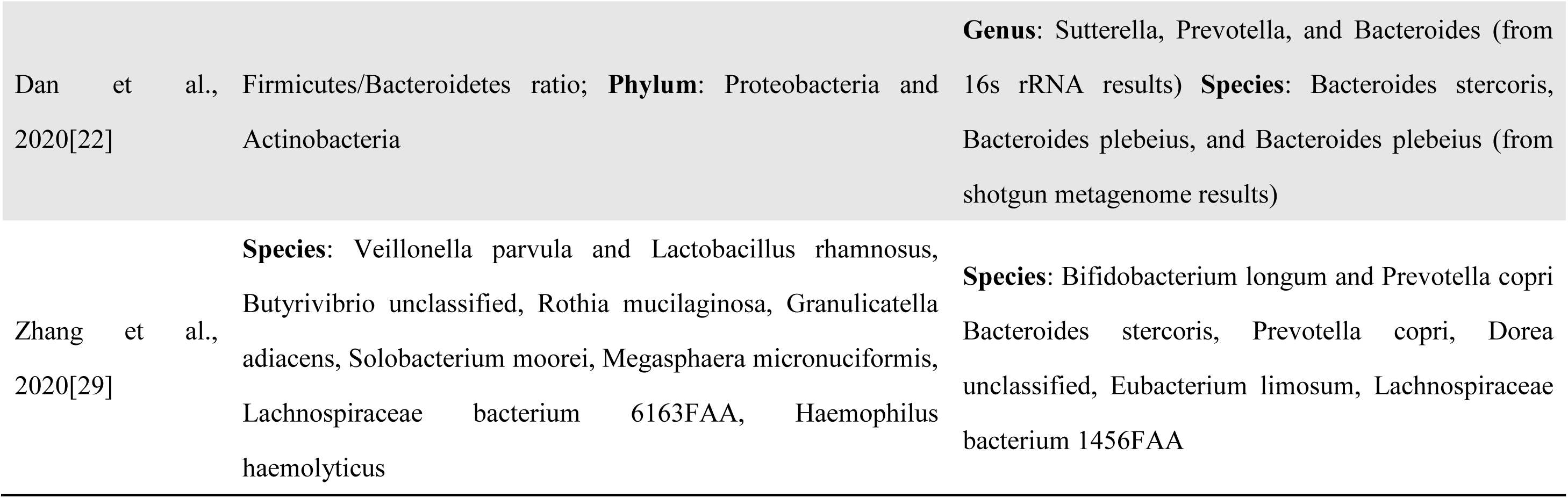
Summary of relevant literature in the past 5 years – results. **Notes:** ASD, autistic spectrum disorder; FGID, functional gastrointestinal disorder; GI, gastrointestinal; NT, neurotypical; N.S: No significant difference. If there is no N.S mark in the last two columns of conclusions, the results are statistically significant.

### Functional Capability Analysis of metagenomic sequencing revealed dysbiosis in the ASD group

In order to investigate the contribution of the microbiota to ASD etiology, we explored the functional roles of gut microbiota in the potential etiology of ASD. As shown in Table 4, pathways associated with propanoate (PWY-821), methyl phosphonate (PWY-7399), cholate (7ALPHADEHYDROX-PWY), sulfur amino acid (PWY-821) and isopropanol (PWY-6876) metabolism were significantly elevated in ASD. In contrast, pathways associated with lipid (PWY-6467), peptidoglycan (PGN) (PWY-6471), glutamate (P162-PWY), preQ0 (PWY-6703), NAD/NADP - NADH/NADPH interconversion (yeast) (PWY-7245, PWY-7268), pyrimidine (PWY-7208, PWY-7197, PWY0-166) and purine (PWY-5695) were significantly decreased in ASD. Additionally, certain energy production-associated pathways (P124-PWY, PWY-5464) were also dysregulated between ASD and TD.

**Table 3.**
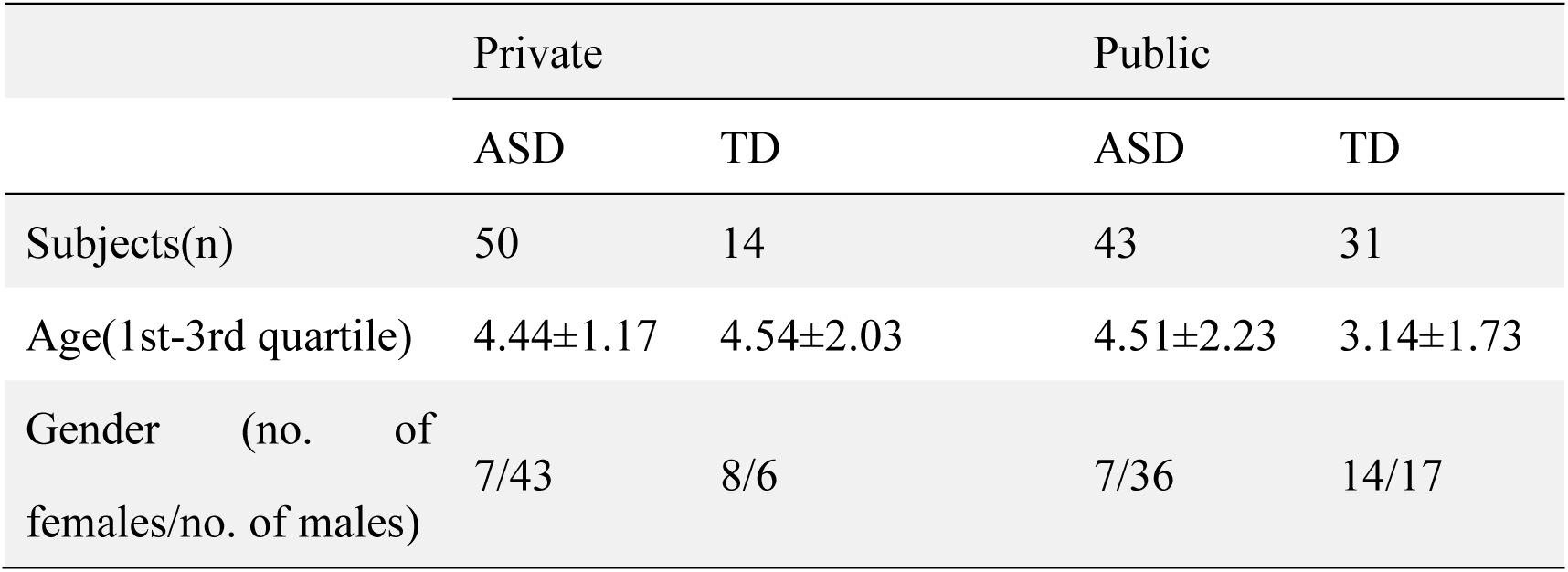
Basic information about the cohort in this study. **Notes:** Age was given as the mean ± standard. ASD, autism spectrum disorder; TD, typical development; MR, mental retardation; SD, speech delay; ADHD, attention deficit hyperactivity disorder.

|  | Private |  | Public |  |
| --- | --- | --- | --- | --- |
|  | ASD | TD | ASD | TD |
| Subjects(n) | 50 | 14 | 43 | 31 |
| Age(1st-3rd quartile) | 4.44±1.17 | 4.54±2.03 | 4.51±2.23 | 3.14±1.73 |
| Gender (no. of females/no. of males) | 7/43 | 8/6 | 7/36 | 14/17 |

**Table 4.**
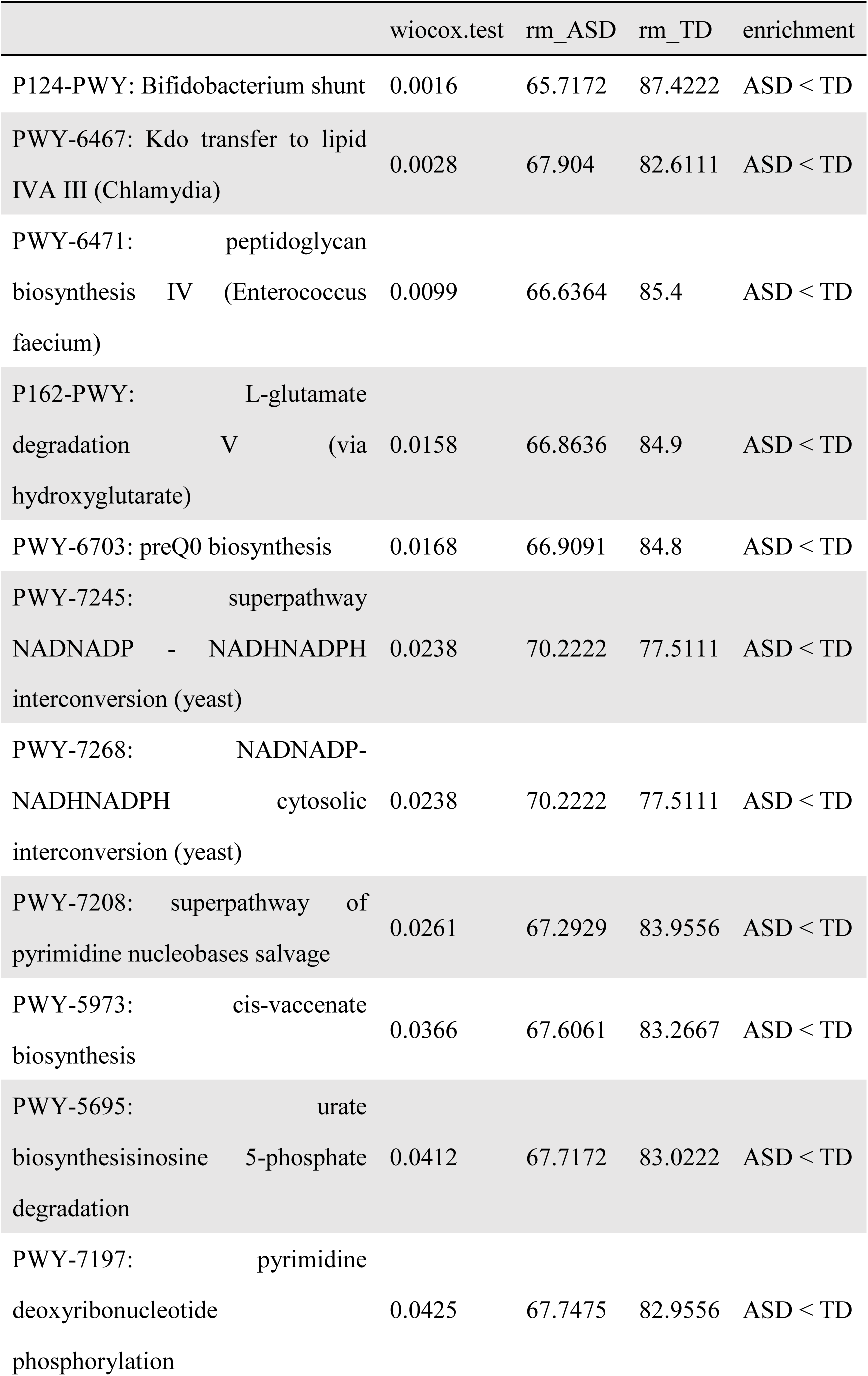
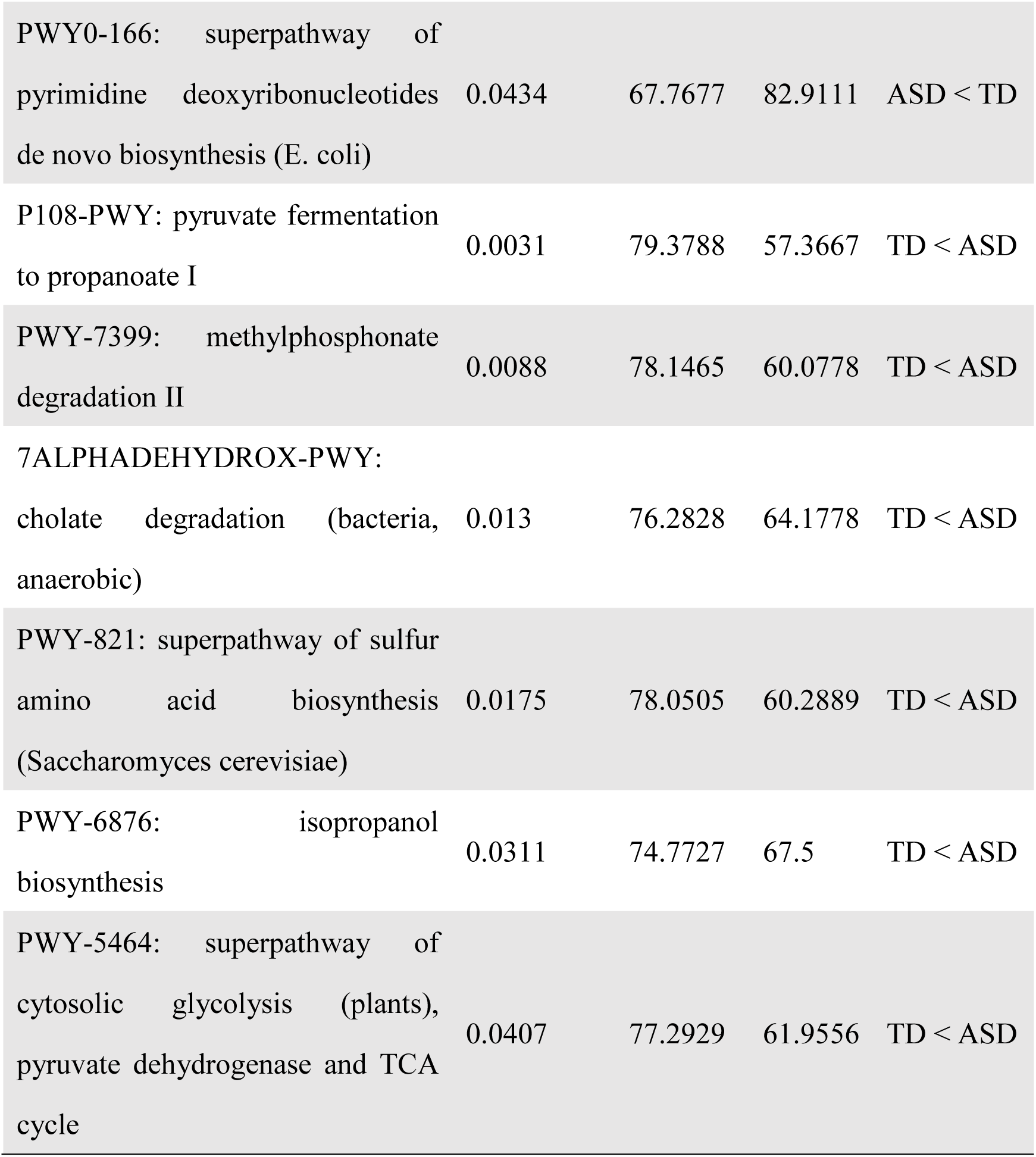
The summary of functional pathways with significant differences. **Notes:** rm: the mean value of rank.

Although cholate degradation pathway was not detected in all ASD samples, there are still many samples with a significantly higher abundance of this pathway than the TD group (Fig. 6A). Notably, cholate degradation in four out of five samples was decreased as behavior improved (Fig. 6B). This pattern was similar to that of *C. scindens* (Fig. 3F). Moreover, *Clostridium* was reported with high bile acid 7alpha-dehydroxylating activity [40, 41]. Obviously, the relative abundance of two species of *Clostridium* were positively correlated with this pathway, especially for *C. scindens* (Fig. 6C). The dysbiosis of this pathway results in elevation of secondary bile acid including lithocholate (LCA) and deoxycholate (DCA) (Fig. 6H). The secondary bile acids produced by bacteria, not only influence signal transport by receptors, such as farnesoid X receptor (FXR), the liver X receptor (LXR), the G protein-coupled receptor TGR5, and the vitamin D receptor [41] but also deplete the membrane of intestinal epithelial cell and mitochondria due to its hydrophobic nature, thereby further influence mitochondrial and epithelial function [42] which is often reported to be involved in the occurrence of ASD [43]. This phenomenon can also be affected by high concentrations of hydrogen sulfide and propionic acid (associated with PWY-821 and PWY-821). Notably, the mitochondrial function of fungi is affected (Table 3, PWY-7245, PWY-7268) which might be the signal of mitochondrial damage which may extend to intestinal epithelial tissue even the brain. The above results indicate the important role of *Clostridium* and the bile acid metabolism pathway in which it participates in autism etiology. Meanwhile, pyrimidine (PWY-7208, PWY-7197, PWY0-166) and purine (PWY-5695) associated pathways were consistently increased in ASD (Fig. 6D-G). Given the crucial roles of purines, pyrimidines, and their derivatives in brain development, alterations in their synthesis, catabolism, and concentrations may lead to significant functional consequences. The increase in excess purine metabolites is regarded as a dangerous cellular metabolic reaction, involving damage to mitochondrial function [44].

**Figure 6.**
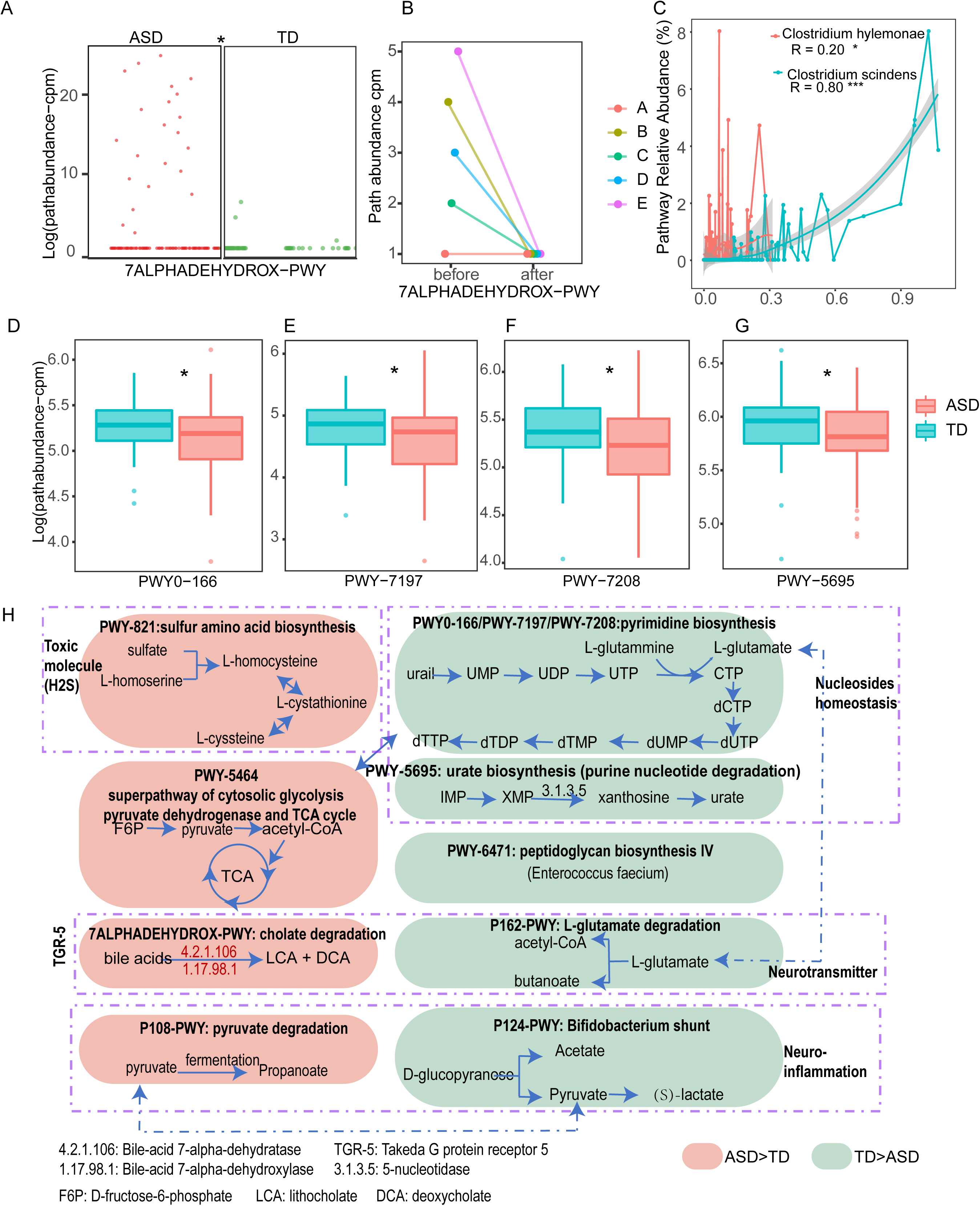
Distinct dysbiosis in functional pathway between ASD and TD. (A) The distribution of abundance of 7ALPHADEHYDROX-PWY: cholate degradation between ASD and TD. (B) Variation of relative abundance of 7ALPHADEHYDROX-PWY of the 5 follow-up ASD children. (C) Correlation between two species (*C. hylemonae, C. scindens*) and the relative abundance of 7ALPHADEHYDROX-PWY. (D-G) comparison of metabolic pathways related to pyrimidine and purine between ASD and TD. (H) Summary and association about dysfunctional metabolic pathways. Values that are significantly different by the Wilcoxon rank sum test are indicated by asterisks as follows: *, *p* < 0.05; **, *p* < 0.01; ***, *p* < 0.001.

As shown in Fig. 6H, we speculated that dysregulation of gut microbiota function is involved in various aspects of autism pathogenesis, including toxic effects, nucleoside homeostasis, neurotransmitters, and neuroinflammation.

## Discussion

Our study was a pilot exploration to examine the association between gut microbiota and ASD etiology using a Chinese cohort based on shotgun metagenomics sequencing. Innovatively, we designed a longitudinal study to verify the results we identified. Several pieces of evidence link the colonization of *Clostridium* bacteria with neurological symptoms and/or ASD etiology in humans [35]. Species belonging to *Clostridium* have been shown to produce exotoxins [45] and p-cresol and promote conditions that favor inflammation that may exacerbate autistic symptoms, especially *Clostridium difficile* [46]. In our study, strain-level identification and verification of the follow-up samples not only further confirmed the significant role of *Clostridium spp*. in ASD etiology but also provided precise references for microbiome intervention and treatment.

Fungi, which can influence the bacterial community, constitute the second-largest flora in the intestine. Because of the small proportion, most studies that associated ASD gut microbiota ignored this community. Although there were still a few studies that indicated higher isolation ratio of yeast or dysbiosis fungi taxon in ASD [11, 47, 48], we discovered for the first time the composition of the fungal community appears to exhibit a higher discrimination between ASD and TD compared to that of the bacterial community. Two pieces of evidence demonstrated an increase of *Candida* in ASD [11, 48]. Moreover, dysbiosis of *C. albicans* can not only lead to long-term alterations in specific bacteria [49], but also trigger innate immune response [50]. Importantly, elevated antibodies against *C. albicans* have been found in individuals with ASD, and mice infected with *C. albicans* display mild memory impairment and trigger an inflammatory response [51, 52]. Our results not only further confirmed the potential role of gut *Candida* dysbiosis in ASD etiology but also narrowed it down to specific species. It is also worth noting that *CX3CR1* and *CLEC7A*, encoding CX3CR1 and Dectin-1 protein by leukocytes that play a decisive role in regulating fungi, are mutated in some ASD patients according to the literature [53–56]. Meanwhile, fungal wall-deficient L-form variants were found in the blood of ASD individuals and the level of mycotoxin in the urine and plasma of ASD patients is higher than that of normal children [57–59]. Therefore, we speculated that the specific genetic variations associated with ASD can regulate the intestinal fungi community by which participation in the pathogenesis of ASD through mycotoxins and immune inflammatory responses (Fig. 7a).

**Figure 7.**
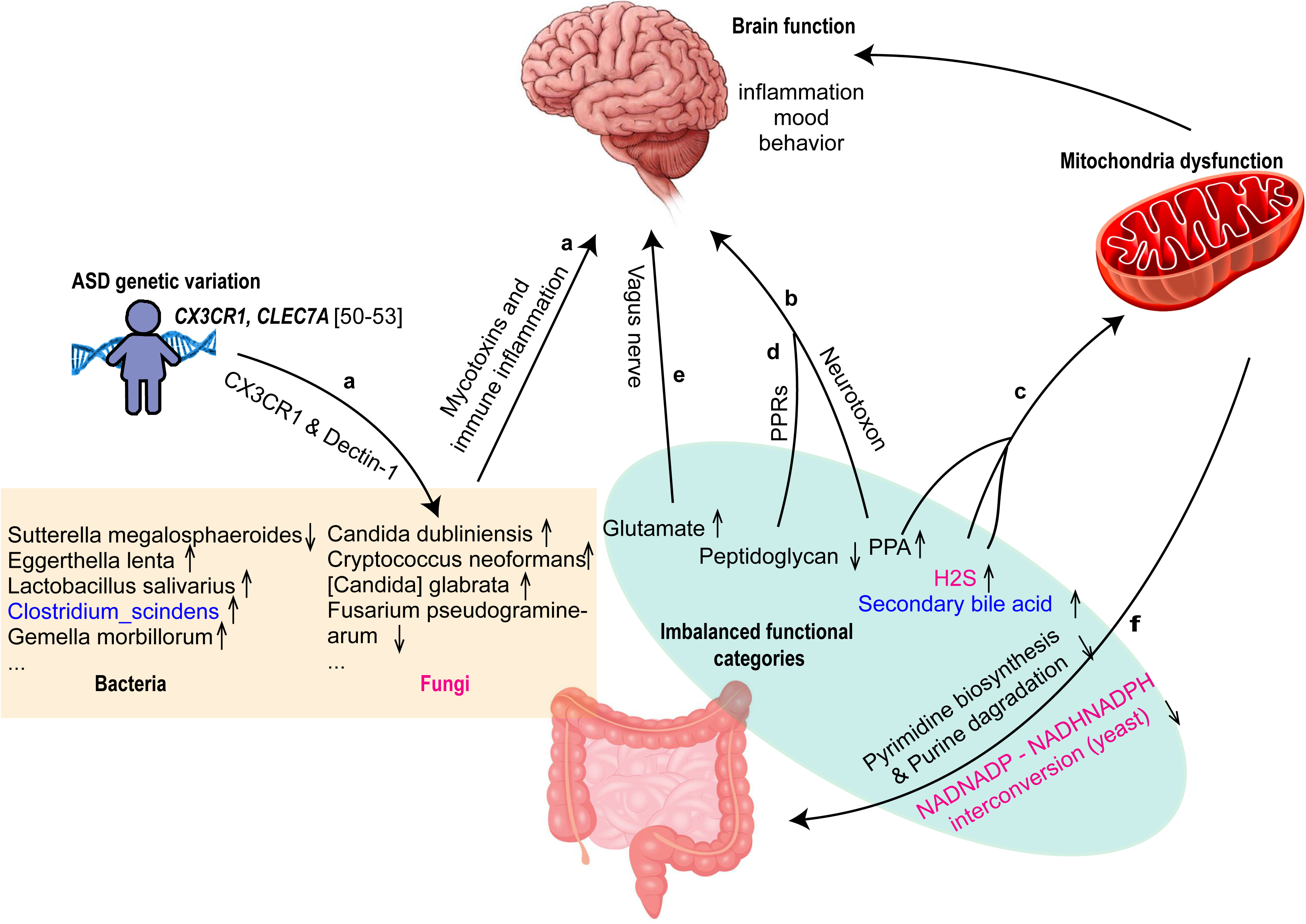
Schematic diagram of the gut microbiota functional pathways involved in the pathogenesis of ASD. **a)** Specific ASD genetic variation can regulate the intestinal fungi community through the protein it encodes thus participate in the pathogenesis of ASD through mycotoxins and immune inflammatory response. **b)** High concentrations of PPA could result in behavioral disorders through neurotoxic characteristic. **c)** The accumulation of H2S, as well as secondary bile acids and propanoic acid (PPA), by affecting the stability of membrane structure due to hydrophobic properties and oxidative stress, affected the expression of specific genes in respiration to influence mitochondrial function. **d)** Peptidoglycan fragments from the bacterial cell wall can be translocated to the brain and activate pattern recognition receptors (PRRs), which were widely expressed in the placenta during the perinatal period at specific developmental stages and participate in the regulation of social behavior, anxiety and stress response behavior. **e)** Dysbiosis glutamate as neurotransmitter through vagus nerve to influence excitatory/inhibitory imbalance associated with the pathogenesis of ASD. **f)** The decreased NAD/NADP - NADH/NADPH interconversion (yeast) that occurred in the yeast mitochondrion might be the direct manifestation of mitochondrial dysfunction in the gut. In the rectangle and the ellipse, the dysbiosis taxon and their participating functional pathways use the same font color to indicate consistency.

Based on the imbalance of the intestinal flora in ASD, we also constructed a high-performing classification model using the relative abundance of species. The top 30 species, selected based on their larger contribution rates in the model, demonstrated complementary integration of results for ASD risk assessment. Moreover, from this list of top 30 species, we also focused on *Bifidobacterium* and *Lactobacillus*. The species distribution of these two genera in individual samples showed a high level of individual heterogeneity, explaining the inconsistency of statistical results across different studies and highlighting the need for more accurate and personalized probiotic interventions for ASD.

Changes in the microbiome often result in altered metabolic profiles, impacting the availability and diversity of nutrients and microbial metabolites [60, 61]. Indeed, metabolic analyses of serum, feces, and urine from ASD subjects have uncovered differences in various molecules compared to TD individuals, with many dysregulated compounds being of microbial origin [18, 19, 23, 62, 63]. Propionate (synonym as propionic acid, PPA) is one of the major neurotoxic SCFA [64]. Elevated concentrations of PPA have been linked to behavioral disorders in various rodent studies (Fig. 7b) [65, 66]. The increased abundance of cholate (primary bile acid) degradation (7ALPHADEHYDROX-PWY) identified in our results leads to the accumulation of secondary bile acid (DCA, deoxycholic; LCA, acid lithocholic acid). We also identified specific strains, such as *C. scindens GCF000154505,* involved in this pathway, and observed a decrease in the abundance of these strains and pathways as the patients’ behavior improved. Studies using high-fat diet mouse models have shown that the hydrophobic properties of deoxycholic acid (DCA) can compromise the integrity of intestinal epithelial cells by affecting the membrane structure stability [67]. Bile acids have also been reported to stimulate the release of 5-HT, a neurotransmitter in CNS, from enterochromaffin (EC) cells [68] which are the most abundant cell type among the enteroendocrine cells throughout the entire GI tract. Patients with Alzheimer’s disease exhibited elevated levels of secondary bile acids in both serum and brain samples, a finding strongly correlated with cognitive decline [69]. Building upon the aforementioned findings in literature that strongly demonstrate the potential roles of bile acids in nervous system diseases, our results provide further insights into the origin of elevated bile acid levels in ASD patients. We employed follow-up samples to validate our findings. However, the causal relationship between the increased abundance of the strain associated with this pathway and its implications still requires thorough investigation by experts in relevant fields.

Growing studies have suggested the importance of mitochondrial dysfunction in the etiology of ASD with both congenital genetic defects in mitochondria and acquired damage caused by environmental factors in ASD [70]. The accumulation of H2S, along with DCA and PPA, influenced by their hydrophobic properties and oxidative stress, can impact membrane stability, [71], ultimately affecting the expression of specific genes involved in respiration and mitochondrial function (Fig. 7c) [72]. Our study speculated that the predicted accumulation of propanoate, bile acids, and H2S based on metagenome data led to mitochondrial dysfunction within the context of the gut microbiota. Meanwhile, we also elucidated the direct consequences of this dysfunction, including the reduction in yeast respiration, as well as subsequent effects, such as compromised pyrimidine biosynthesis and impaired purine degradation, subsequently influencing the gut and brain. (Fig. 7c, f).

Peptidoglycan fragments from the bacterial cell wall can be translocated to the brain and activate pattern recognition receptors (PRRs), which were widely expressed in the placenta during the perinatal period at specific developmental stages and participate in the regulation of social behavior, anxiety and stress response behavior (Fig. 7d) [73]. We also found that glutamate degradation pathway was significantly decreased in the guts of children with ASD compared to controls. This may result in the accumulation of glutamate in ASD gut (Fig. 7e). Increased levels of amino acid transporters in the intestines of the mouse model of ASD were found to contribute to the high level of serum glutamine and the increased E/I ratio in the brain [8]. Accumulation of glutamate in ASD gut may be the resource of high level of serum glutamine.

Taken together, at functional pathway level, we assume that ASD children’s intestinal microbes influence mitochondrial damage, endogenous immune reactions, and neurotoxicity through specific metabolites. These impacts can disrupt the normal function of intestinal tissue and potentially affect the brain, including the regulation of the vagus nerve neurotransmitter. Subsequently, these harmful effects may contribute to metabolic disorders of certain substances within gut microbes (Fig. 7).

Although we did find significant differences in the composition and functionality of gut microbiota between ASD and TD, this is not sufficient evidence to directly prove that these differences can directly contribute to the etiopathogenesis of ASD. Diet is known to directly alter the composition of gut microbiota [74, 75], therefore, homogenous diets often observed in ASD children could result in less intra-individual but more inter-individual variety in gut microbiota as well. In a recent autism stool metagenomics study, genetic measures and restricted dietary preferences seemed to result in less diverse gut microbiota and additionally, looser stool consistency [76]. In our results, gut microbiota appears to be a side contributive factor to exacerbate gastrointestinal and behavioral symptoms in ASD. Our longitudinal study directly indicates the relationship between gut microbiota and the behavioral symptoms of autism.

There are still limitations in the present study. The high individual variability between samples and small size of follow-up samples only to a certain extent to verify the results.

Further cohorts and broader perspectives are needed, including metagenomic, meta-transcriptomic, and metabolomic profiles, as well as host genome information.

## Conclusion

In conclusion, empirical data and computational analyses highlight the significant involvement of gut microbiota composition and intestinal dysbiosis in ASD etiology. Specifically, we emphasize the substantial contribution of fungi to ASD etiology and elucidate the intricate influence of gut microbiota on neurotransmitters, the innate immune system, and mitochondrial dysfunction, collectively shaping ASD pathogenesis. Additionally, our findings uncover both direct and indirect consequences of mitochondrial damage. These insights provide a comprehensive perspective for ASD intervention, emphasizing innovative approaches informed by microbiota-based interventions such as probiotics, fecal microbiota transplantation, or metabolites, which present timely and promising strategies to address the enduring challenges of ASD.

## Materials and methods

### Study participants and samples handling and collection

In the present study, we enrolled 50 ASD children from the Center of Wucailu Child Behavioral Intervention and 14 TD children from nursery and primary schools in Beijing, China. Table 1 shows the general characteristics of participants. The diagnosis of ASD was based on After Autism Diagnostic Observation Schedule (ADOS) evaluation, which was the current ‘gold standard’ diagnostic tool. All stool samples of enrollment of ASD children performed shotgun metagenome sequencing.

Considering the limitation of sample size of our enrolled cohort, we combined age and ethnic matched public metagenome dataset that consisted of 43 ASD children and 31 TD children [23]. Overall, there is a total of 92 children with ASD and 42 TD children in the combined cohort and all downstream analyses, derived from the sequencing raw data, were conducted using a uniform pipeline.

Following the enrollment of ASD children, we conducted a year-and-a-half-long follow-up during which these children received Applied Behavior Analysis (ABA) interventions. Before and after intervention, we performed individualized Psychoeducational Profile, Third Edition (PEP-3) assessments for each ASD child. PEP-3 Assesses skills and behaviors of children with autism.Five children who exhibited notable improvements performed a second round of stool sample collection for in-depth validation analysis.

All the participants did not receive any antibiotic treatment, probiotics, prebiotics or any other medical treatment that could influence the intestinal microbiota during at least 2 months before they were enrolled in the study.

Fecal specimens were collected in the homes of the participants by their parents. Immediately deep freezing was required to preserve the specimens, and then fecal specimens were shipped to the laboratory where each specimen was frozen at −80 °C until DNA extraction.

### Library preparation and Illumina sequencing

Libraries for DNA were prepared using the TruSeq method and sequenced as meta library(350bp) each multiplexed through Illumina HiSeqXten machines using the 2×150 bp paired-end read protocol at the Novogene Co., Ltd. Reads were preprocessed based on quality scores using the FASTX suite with default parameters. an average of 25,712,760 (range 17,265,809 −40,069,242) DNA reads and 34365610 (range 29394760 - 41803304) RNA reads per sample (Table S1) were obtained. The amount of sequence collected for each sample is summarized in Table S1.

### Microbiome bioinformatics

Bowtie2 v2.3.2 [77] was used to map reads to the human genome for decontamination before subsequent analysis. Taxonomic and functional profiling was conducted using MetaPhlAn2 [24], kraken2 [25] and HUMAnN2 [26]. Taxonomic profiling of the metagenomic samples was performed using MetaPhlAn2 v2.6.0, which uses a library of clade-specific markers to provide pan-microbial (bacterial, archaeal, viral and eukaryotic) quantification at the species level. MetaPhlAn2 was run using default settings. Functional profiling was performed with HUMAnN2. For an input metagenome, HUMAnN2 constructed a sample-specific reference database by concatenating and indexing the pangenomes of species detected in the sample by MetaPhlAn2 (pangenomes were pre-clustered, pre-annotated catalogues of open reading frames found across isolate genomes from a given species). HUMAnN2 then mapped sample reads against this database to quantify gene presence and abundance in a species-stratified manner, with unmapped reads further used in a translated search against UniRef90 [78] to include taxonomically unclassified but functionally distinct gene family abundances. Finally, for community-total, species-stratified, and unclassified gene family abundance, HUMAnN2 reconstructed metabolic pathway abundance based on the subset of gene families annotated to metabolic reactions (based on reaction and pathway definitions from MetaCyc. Enzyme (level-4 Enzyme Commission (EC) categories) abundances were further computed by summing the abundances of individual gene families annotated to each EC number based on UniRef90-EC annotations from UniProt [79].

### Statistical analysis

Continuous variables were expressed as mean ± standard deviation, and comparisons between the ASD and neurotypical groups were conducted using the Wilcoxon rank-sum test implemented in the coin package in R (http://cran.r-project.org/). To identify features differentially represented between any two groups, we also employed LEfSe (linear discriminant analysis effect size) [31], an algorithm utilizing linear discriminant analysis (LDA) to estimate the effect size of taxa that are differentially represented between two compared groups. The relative abundance of each taxon with adjusted *P* < 0.05 and LDA >2 as significant. At other conditions, p values lower than 0.05 were accepted as significant. The gut microbiota Shannon diversity analysis was performed using Vegans in the R package at species-level data. To facilitate interpretation of the results, non-metric multidimensional scaling (NMDS) was performed based on flower composition Bray-Curtis distances using the metaMDS function of the vegan package in R. The permutational multivariate analysis of variance (PERMANOVA) Analysis (or Adonis test) was conducted using the adonis function of the vegan package in R with 999 permutations. Pearson correlation coefficient was conducted using cor. test function in R.

### Random forest model

The construction of the random forest model utilized the randomForest R package (https://cran.r-project.org/web/packages/randomForest/index.html). The model was trained using 75% of the samples via fourfold cross-validation and tested using all samples, with bootstrapping performed 1000 times. The diagnostic capacity of the model in discriminating ASD individuals from control subjects was assessed using the AUC (area under the ROC curve) via the R package pROC (https://cran.r-project.org/web/packages/pROC/index.html). The mean contributions of each species were calculated to evaluate their differences between the ASD and TD groups.

## Data availability

The raw sequence data reported in this paper have been deposited in the Genome Sequence Archive [80] in National Genomics Data Center [81], Beijing Institute of Genomics and China National Center for Bioinformation [82], Chinese Academy of Sciences (GSA: CRA007507) that are publicly accessible at https://ngdc.cncb.ac.cn/gsa.

## Competing interests

The authors declare no competing interests.

## Acknowledgments

This work was supported by the Strategic Priority Research Program of the Chinese Academy of Sciences [XDB38030200]; Genomics Data Center Operation and Maintenance of Chinese Academy of Sciences [CAS-WX2022SDC-XK05]; The Alliance of International Science Organizations [ANSO-PA-2023-07]

---

**Figure S1.**
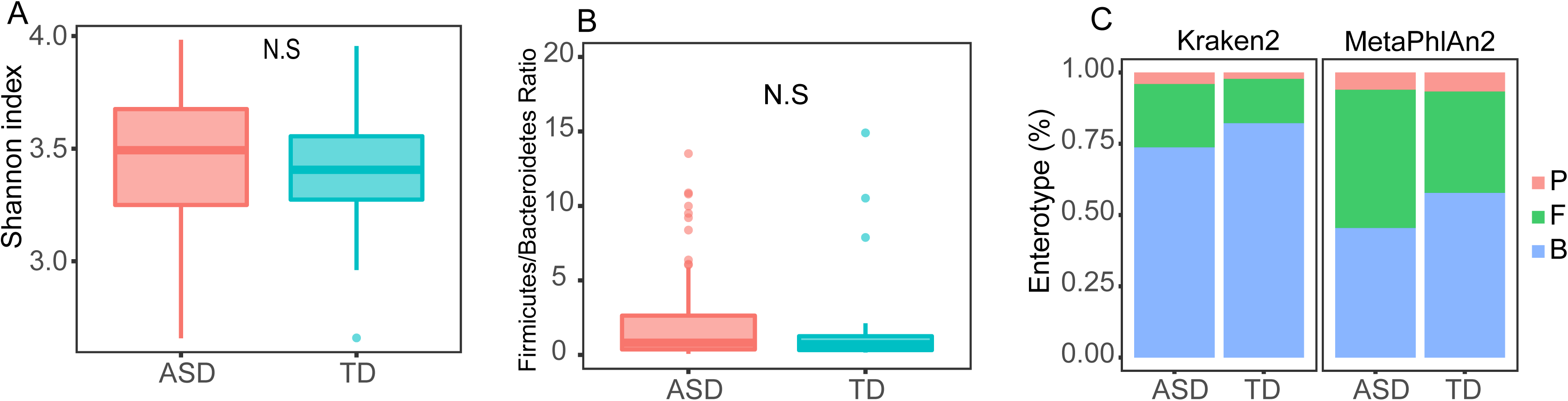
(A) Shannon diversities were independently calculated for the ASD and TD groups. (B) Comparison on Firmicutes/Bacteroidetes ratio between ASD and TD. (C) Enterotypes of the ASD and TD groups based on genus level quantification that were evaluated by Kraken2 and MetaPhlAn2.

**Figure S2.**
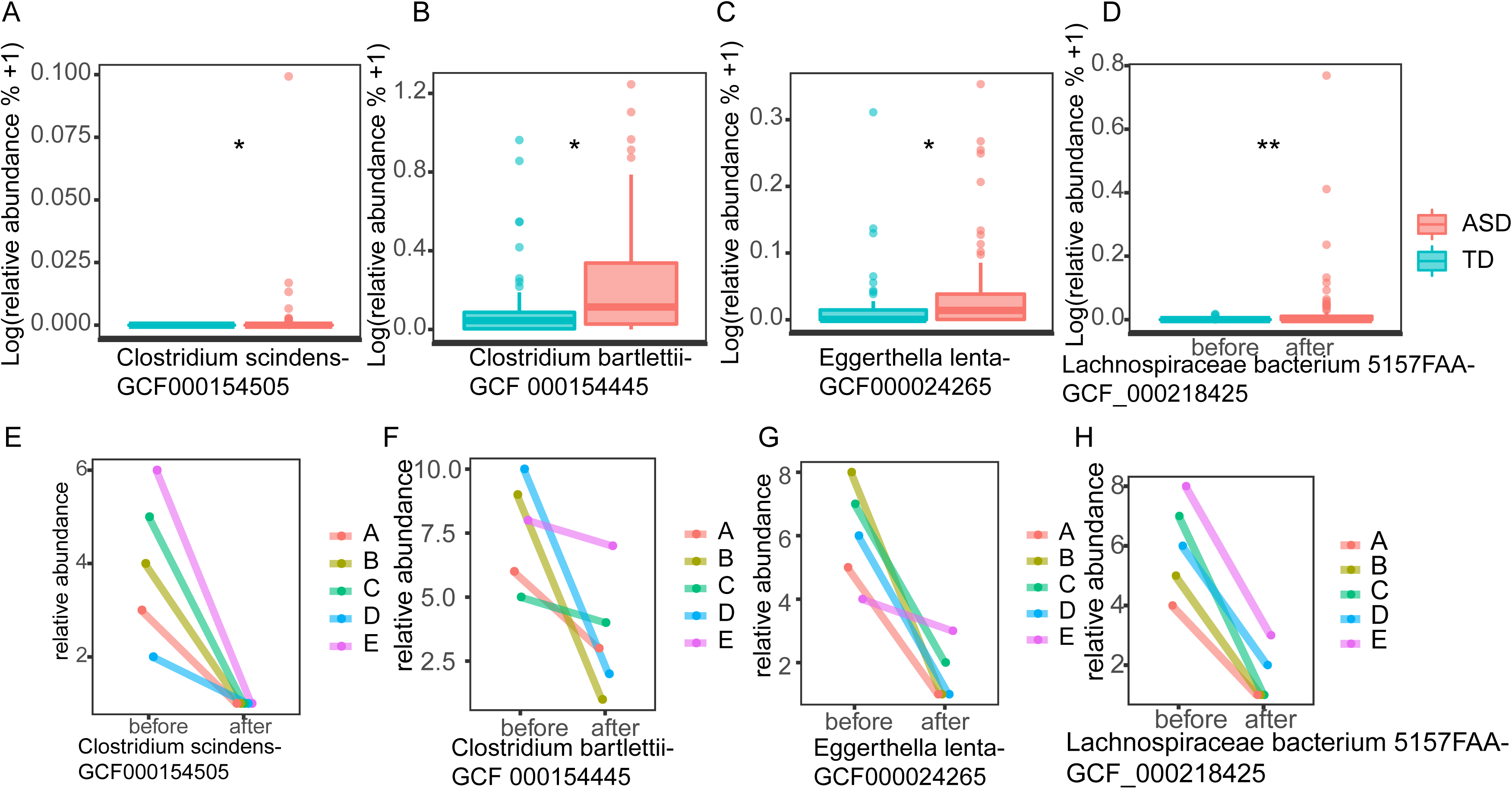
(A-D) Comparison of the relative abundance of 4 strains between ASD and TD. (E-H) Variation of relative abundance of 4 strains in 5 follow-up ASD children. Values that are significantly different by the Wilcoxon rank sum test are indicated by asterisks as follows: *, adj.*p* < 0.05; **, adj.*p* < 0.01; ***, adj.*p* < 0.001.

**Figure S3.**
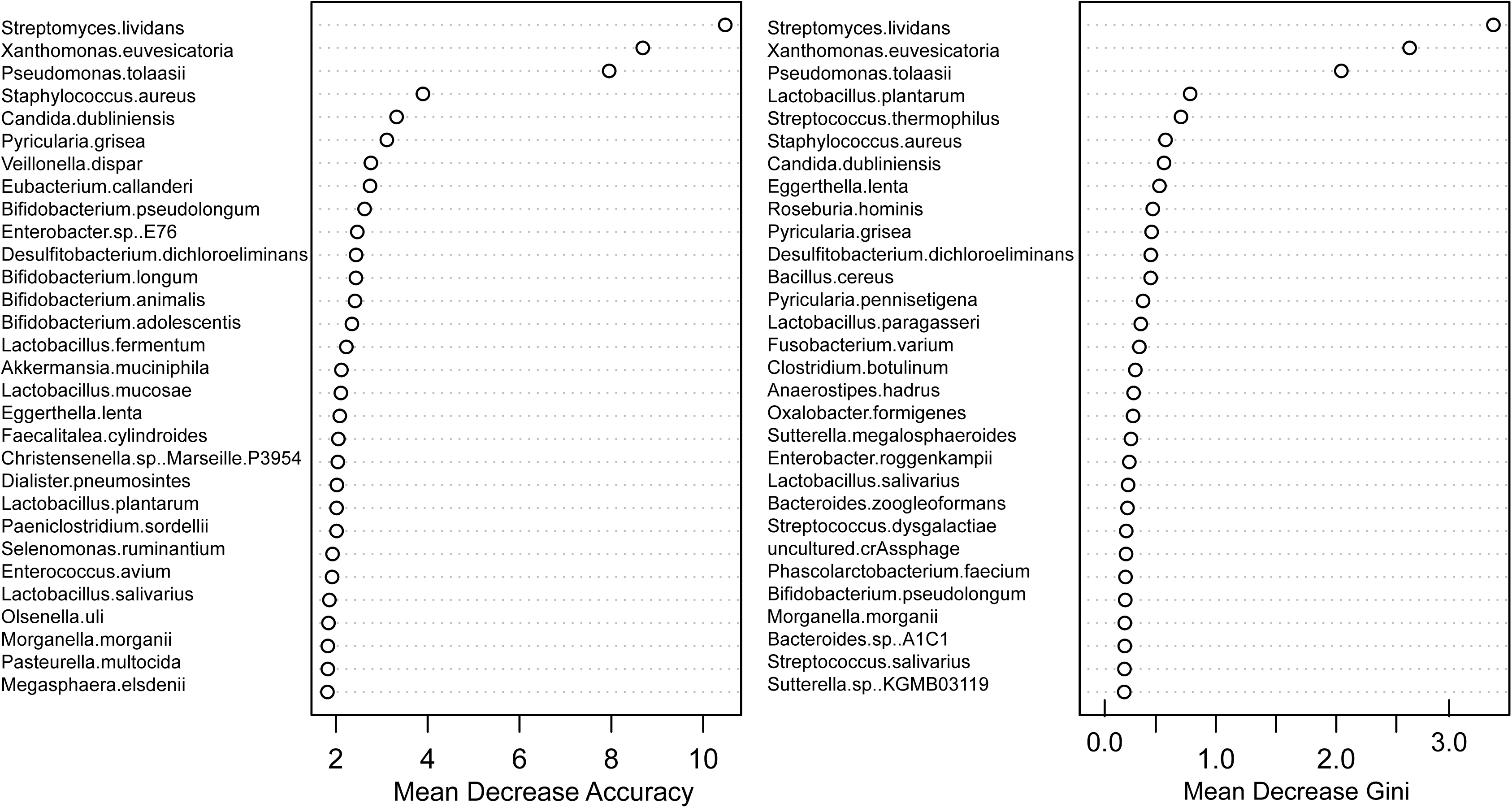
Top 30 variable importance based on Mean Decrease Accuracy and Mean Decrease Gini from random forest model.

## Notes

### Competing Interest Statement

The authors have declared no competing interest.

